# Fitness effects of *cis*-regulatory variants in the *Saccharomyces cerevisiae TDH3* promoter

**DOI:** 10.1101/154039

**Authors:** Fabien Duveau, William Toubiana, Patricia J. Wittkopp

## Abstract

Variation in gene expression is widespread within and between species, but fitness consequences of this variation are generally unknown. Here we use mutations in the *Saccharomyces cerevisiae TDH3* promoter to assess how changes in *TDH3* expression affect cell growth. From these data, we predict the fitness consequences of *de novo* mutations and natural polymorphisms in the *TDH3* promoter. Nearly all mutations and polymorphisms in the *TDH3* promoter were found to have no significant effect on fitness in the environment assayed, suggesting that the wild type allele of this promoter is robust to the effects of most new *cis*-regulatory mutations.

## Main text

Genomic studies have identified extensive variation in gene expression within and between species (reviewed in Ranz and Machado 2006; Alvarez et al. 2015). Given the key role that gene expression plays in regulating development and physiology, many of these changes in expression are expected to affect higher order phenotypes, which may in turn affect fitness (reviewed in Gilad et al. 2006; Wray 2007; Carroll 2008; Fay and Wittkopp 2008; Wittkopp and Kalay 2012). Despite these often-stated expectations, only a handful of studies have directly measured the fitness effects of changes in gene expression (Dykhuizen et al. 1987; Perfeito et al. 2011; Rest et al. 2013; Keren et al. 2016; Rich et al. 2016; Bergen et al. 2016), and most of these studies have used heterologous or synthetic promoters to alter gene expression level. It therefore remains largely unknown how differences in gene expression induced by new mutations in regulatory sequences relate to changes in organismal fitness. Here we introduced mutations in the promoter of the *TDH3* gene in *Saccharomyces cerevisiae* to alter *TDH3* expression levels and determine the impact of these expression changes on fitness during clonal growth in a rich medium containing glucose, which is the preferred carbon source for *S. cerevisiae* (Gancedo 1998). We then use this information to predict the fitness effects of new mutations and natural polymorphisms within the *TDH3* promoter that were previously characterized for their effects on expression (Metzger et al. 2015).

*TDH3* encodes a glyceraldehyde-3-phosphate dehydrogenase best known for its role in central metabolism (McAlister and Holland 1985) but also implicated in silencing of Sir2-dependent genes in telomeric regions (Ringel et al. 2013) and observed among antimicrobial peptides secreted by *S. cerevisiae* during alcoholic fermentation (Branco et al. 2014). Prior work has shown that eliminating *TDH3* decreased fitness in rich media (YPD) by ~4% (Deutschbauer et al. 2005) and overexpressing *TDH3* inhibited expression of telomeric genes (Ringel et al. 2013), suggesting that fitness (*i.e*. population growth rate) should be sensitive to *TDH3* expression level. To determine the fitness effects of altered *TDH3* expression levels, we selected eight G->A or C->T point mutations in the *TDH3* promoter (*P_TDH3_*) that caused a broad range of changes in expression of a yellow fluorescent protein (YFP) reporter gene (Figure 1A; Metzger et al. 2015) and introduced them into the native *TDH3* locus (Figure 1B). To expand the range of expression changes assayed, we also constructed and analyzed three strains with duplications of the reporter gene separated by a *URA3* marker as well as a strain with only the *URA3* marker inserted downstream of the wild type *P_TDH3_-YFP* reporter gene to serve as a control for the duplication strains (Figure 1A). One of these duplications contained two copies of the wild type *P_TDH3_* allele and the other two duplications each carried a single mutation in both copies of the promoter (Figure 1A). A matching set of strains containing duplications of the *TDH3* gene was also constructed and used to measure fitness (Figure 1B). We used the parent of the reporter gene strains (BY4724), which lacks the *P_TDH3_-YFP* reporter gene (labeled “Deletion” in Figure 1A), to determine the level of auto-fluorescence in the absence of YFP, and we deleted the promoter and coding sequence of the native *TDH3* gene (labeled “Deletion” in Figure 1B) to quantify fitness when *TDH3* was not expressed. Prior work has shown that the effects of mutations in the *P_TDH3_-YFP* reporter gene on fluorescence levels are nearly perfectly correlated (R^2^ > 0.99) with the effects of the same mutations at the native *TDH3* locus measured using a *TDH3::YFP* fusion protein (Metzger et al. 2016).

**Figure 1.**
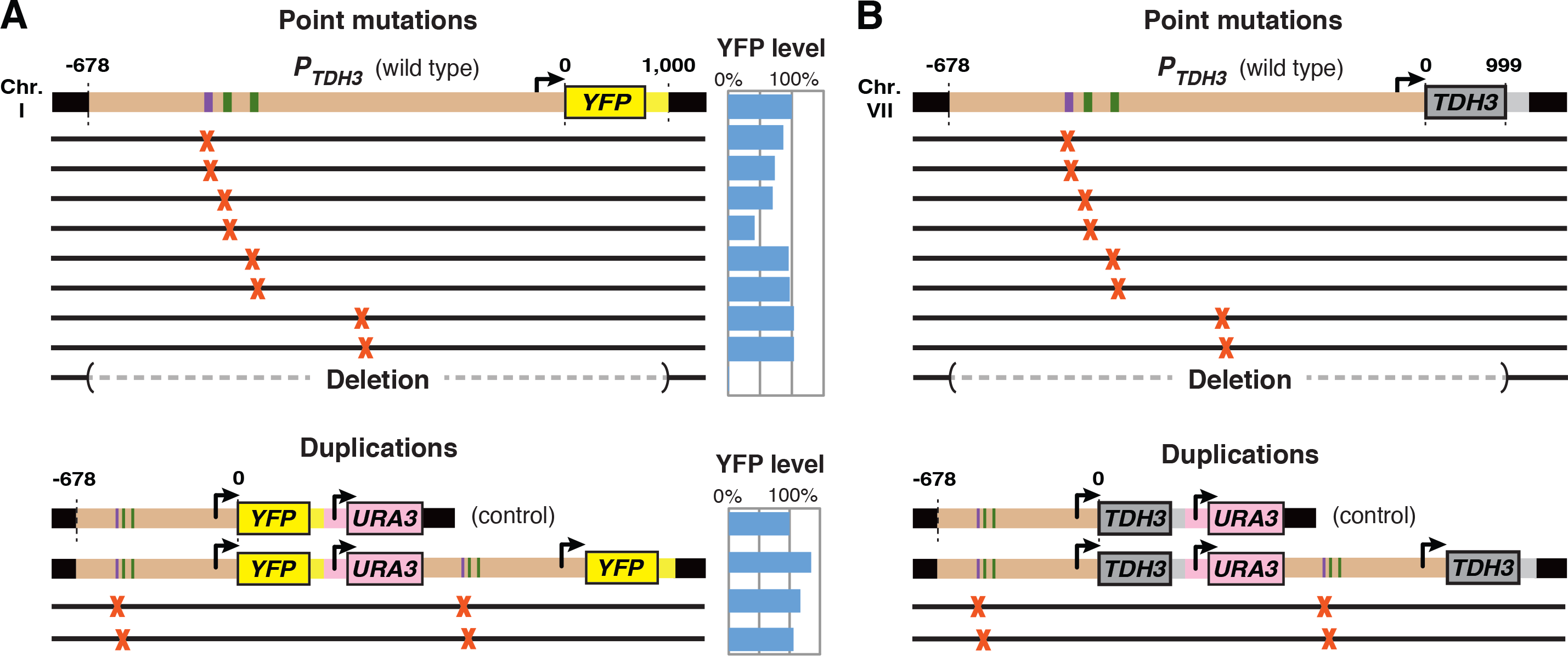
Genomic constructs used to alter *TDH3* expression and measure effects on fitness. Schematics show the 14 genomic constructs used to quantify the effects of different alleles of the *TDH3* promoter (*P_TDH3_*) on (A) expression level and (B) fitness. (A,B) Wild type alleles of *P_TDH3_* are shown in brown with transcription factor binding sites for RAP1 (purple) and GCR1 (green) indicated. Arrows show transcription start sites, and thick black lines represent surrounding genomic sequence. Thinner black lines represent genomic constructs with point mutations (G->A or C->T) in *P_TDH3_* at sites indicated with red “X”s. Constructs shown in (A) have these alleles fused to *YFP* coding sequence and the *CYC1* transcription terminator and were inserted into chromosome I at position 199,270. From top to bottom, these schematics represent the wild type *P_TDH3_-YFP* allele, eight alleles with single point mutation in *P_TDH3_*, the parental BY4724 strain lacking *P_TDH3_-YFP*, the control for duplication alleles with a *URA3* marker inserted downstream of the wild type *P_TDH3_-YFP* allele, the duplication of the wild type *P_TDH3_-YFP* separated by the *URA3* marker, and two duplication alleles that each has a point mutation in both copies of *P_TDH3_*. Blue bars to the right of these 14 schematics show the average fluorescence level of each construct relative to the wild type allele. Constructs shown in (B) have these *P_TDH3_* alleles inserted at the native *TDH3* locus on chromosome VII. From top to bottom, they represent the wild type *TDH3* gene, eight alleles with single point mutations in *P_TDH3_*, the null allele created by a deletion spanning the 678 bp promoter and the entire 999 bp *TDH3* coding sequence, the control for duplication alleles with a *URA3* marker inserted downstream of the native *TDH3* gene, the duplication of the wild type *TDH3* gene (promoter, coding sequence, and transcription terminator), and two *TDH3* duplication alleles that each contains a single point mutation in both copies of the *P_TDH3_* promoter.

The expression level driven by these 14 *P_TDH3_* alleles was estimated using the fluorescence level of the reporter gene strains measured by flow cytometry after growth in a rich medium (YPD). Relative median fluorescence of each genotype (shown in Figure 1A) was determined by averaging median fluorescence from at least four replicate populations and dividing it by the average fluorescence of the corresponding wild type allele. To quantify the impact of changing *TDH3* expression on fitness, each of the 14 *TDH3* strains (Figure 1B) was co-cultured with a common competitor strain in the same growth conditions (Figure 2A). Abundance of each strain relative to the competitor was quantified in at least seven replicates at four time points during ~21 generations of growth (Figure 2B), and then relative fitness was calculated for each genotype by dividing the average competitive fitness across replicates by the average competitive fitness of the genotype carrying the corresponding wild type allele of *P_TDH3_* (Figure 2A).

**Figure 2.**
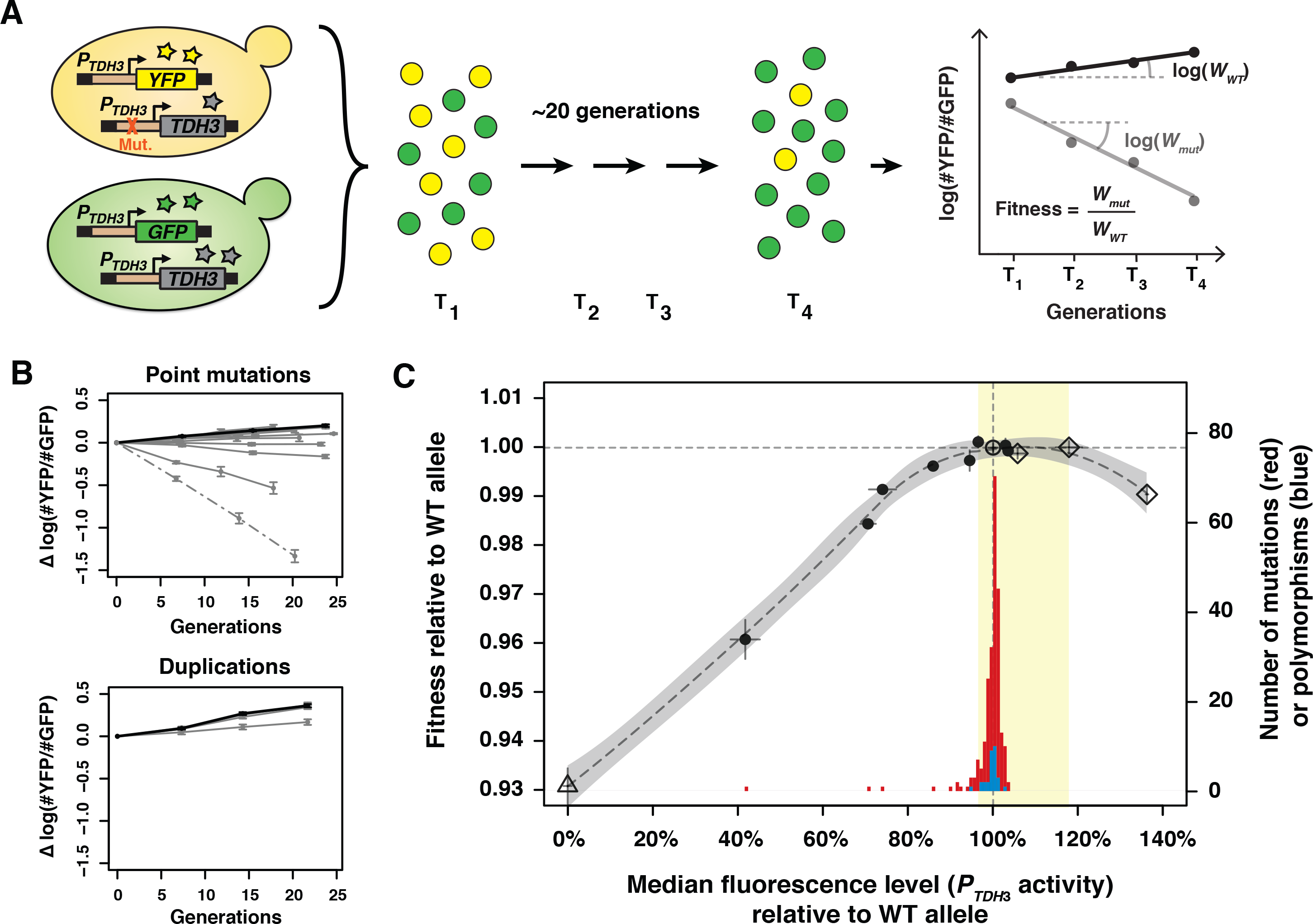
Fitness effects of *cis*-regulatory mutations and polymorphisms in the *TDH3* promoter. (A) An overview of the competition assay used to quantify relative fitness is shown. Each of the 14 *TDH3* alleles shown in Figure 1B was introduced into a strain carrying wild type *P_TDH3_-YFP* inserted at the *HO* locus, which as used to mark cells, not to measure expression. The resulting strains were competed individually against a strain wild type for *TDH3* that was marked with a green fluorescent protein (*P_TDH3_-GFP* reporter gene at *HO*. Each of the 14 [YFP+] strains was mixed with the [GFP+] strain in at least seven replicates and grown at log-phase in rich medium (YPD) for 30 hours (~20 generations). The frequency of [YFP+] and [GFP+] cells was quantified by flow cytometry at four time points, once every 10 hours, at which points cell cultures were diluted to fresh medium. Competitive fitness (*W*) was determined from changes in the frequency of [YFP+] and [GFP+] cells over time. The logarithm of the relative frequency of [YFP+] and [GFP+] cells was regressed on the number of generations of growth and the exponential of the regression coefficient corresponded to the competitive fitness of the [YFP+] strain. To account for the small difference in growth rate between the [YFP+] strain carrying the wild type allele of *TDH3* and the [GFP+] competitor strain, relative fitness was calculated by dividing the average competitive fitness measured for each mutant strain (*W_mut_*) by the average competitive fitness measured for the strain with the wild type allele of *TDH3* (*W_WT_*). (B) Changes in genotype frequency over time (number of generations) are shown for the eight alleles with single point mutations in *P_TDH3_* (top; plain grey lines), for the deletion of *TDH3* (top; dotted line), for the three alleles with duplication of the entire *TDH3* locus (bottom, plain grey lines) and for the corresponding control strains (black lines). For better visualization, variation in the starting frequency of [YFP+] and [GFP+] cells was removed by subtracting the logarithm of the ratio of [YFP+] and [GFP+] cells measured at the first time point from the ratio measured at each time point (y-axis). Dots indicate the average value of this corrected ratio across all replicate populations for each genotype and at each time point, while error bars show 95% confidence intervals across replicates. Note that the frequency of the wild type [YFP+] strain slightly increased over time (top; black line) and that the control strain with the *URA3* marker (bottom; black line) grew slightly faster than the wild type strain without *URA3* (top; black line), reflecting the small fitness advantage conferred by the *URA3* marker in rich medium. (C) The relationship between relative *P_TDH3_* activity, measured by fluorescence of the *YFP* reporter genes shown in (A), and relative fitness is shown by the black points and the dotted curve. Each point represents the mean of median fluorescence levels (x-axis) among at least four replicate populations and the mean of relative fitness measurements (left y-axis) from at least seven replicate competition assays, with different markers used for the wild type allele (open circle), point mutations (closed circles), null allele (open triangle), and duplication alleles (open diamonds). Error bars show the 95% confidence intervals for both measurements. The dotted curve was defined using a LOESS regression of the fluorescence data on fitness, with the greyed interval around the curve representing the 95% confidence interval of this LOESS regression. The range of *TDH3* expression levels with fitness comparable to the wild-type allele is shown in yellow. Histograms showing the distributions of effects on *P_TDH3_* activity (as measured by fluorescence of the YFP reporter gene) for 235 mutations (red) and 30 polymorphisms (blue) are overlaid on the fitness curve. Each bar represents a bin of 0.8% on the x-axis, and the number of mutants in each bin is shown on the right y-axis. The vertical and horizontal dotted lines show the fluorescence level and fitness conferred by the wild type *TDH3* promoter allele.

These data allowed us to determine the relationship between *TDH3* expression level and fitness upon growth in a rich medium (Figure 2C). We found that a decrease in *TDH3* expression level as small as 6% relative to the wild type level (as measured by YFP fluorescence) significantly reduced fitness by 0.3% (*t*-test, t = −2.08, P = 0.045), whereas a 36% increase in expression level significantly decreased fitness by 1.0% (*t*-test, t = -10.70, P = 2.11 x 10^-7^). The decrease in fitness associated with overexpression of *TDH3*, which is one of the most highly expressed gene in *S. cerevisiae* (Newman et al, 2006), could be due to the energetic cost of protein expression (Kafri et al, 2016, Makanae et al, 2013) or to specific functions of *TDH3* such as an increase in the silencing of telomeric genes (Ringel et al, 2013). No significant changes in fitness were observed for *P_TDH3_* alleles driving expression levels between 96% (*t*-test, t = 1.67, P = 0.10) and 118% (*t*-test, t =0.08, P = 0.94) of the wild type level, indicating a plateau of maximal fitness for expression levels in this range (Figure 2C). This plateau might extend to even higher expression levels given that it is defined by only one overexpression allele with a significant reduction in fitness, and this allele, a duplication of the native *TDH3* gene, showed a 136% increase in fluorescence rather than the doubling reported previously for a duplication of *P_TDH3_* (Kafri et al. 2016), suggesting that the flow cytometer underestimated YFP abundance at high fluorescence levels. Differences in expression noise (variability among genetically identical cells) among *P_TDH3_* alleles might also contribute to differences in fitness among these *P_TDH3_* alleles (Metzger et al. 2015), but the fitness effects of changing expression noise are expected to be much smaller than the fitness effects of changing mean expression level (Keren et al. 2016, Wang and Zhang 2011). The data presented here are insufficient to disentangle the fitness effects of changing expression noise and mean expression level because the *P_TDH3_* alleles examined show strongly correlated effects on these two measures (Supplementary Figure 1).

Using the relationship we observed between changes in *TDH3* expression level and relative fitness, we inferred the fitness effects of 235 G->A or C->T point mutations in *P_TDH3_* previously characterized for their effects on YFP fluorescence in the same environment (Metzger et al, 2015). We found that 216 (92%) of these 235 mutations caused changes in expression that are not predicted to affect fitness in this environment (red distribution in Figure 2C). None of these mutations increased expression more than 4% relative to the wild type allele, and the distribution of mutational effects was concentrated at the lower expression end of the fitness plateau (Figure 2C), suggesting that a mutational bias limits increases in *TDH3* expression from arising in similar environments. To understand how selection has filtered this mutational variation in natural populations, we also examined the effects of 30 unique polymorphisms on *P_TDH3_* described in Metzger et al. (2015). We found that 29 (97%) of these 30 polymorphisms caused expression levels within the observed plateau of maximal fitness (blue distribution in Figure 2C), consistent with the absence of evidence for selection acting on *P_TDH3_* activity level in a glucose-based medium reported previously (Metzger et al. 2015). The effects of polymorphisms were also concentrated near the lower expression end of the fitness plateau (maximum level = 103%), again suggesting that natural variation in the activity of the *TDH3* promoter is constrained by mutational biases. Growth in other environments might change the effects of *P_TDH3_* mutations on gene expression and/or the relationship between *TDH3* expression level and fitness {Keren:2016fx} and should be investigated in future work. Effects of different types of mutations (*e.g*. indels, duplications, aneuploidies) should also be examined to determine how they might change the distribution of mutational effects.

Finding that most *cis*-regulatory mutations and polymorphisms cause changes in *TDH3* expression level within the observed fitness plateau indicates that their effects are largely buffered at the fitness level, conferring robustness in the activity of the wild type *P_TDH3_* allele against new mutations, at least in the environment assayed. Non-linear fitness functions with plateaus located around the wild type level of gene expression have previously been described in *S. cerevisiae* for the *LCB2* gene using an inducible promoter (Rest et al. 2013) as well as for other yeast genes using synthetic promoters (Keren et al. 2016) or interspecific comparisons (Bergen et al. 2016), and are consistent with predictions from metabolic flux control theory (Kacser and Burns 1973). These observations suggest that fitness plateaus might be common for expression-fitness functions and might play an important role in shaping the diversity of gene expression levels observed in natural populations.

## Supplementary Materials

Supplementary Materials include (i) Supplementary Text with the Materials and Methods section and a supplementary figure showing the relationship between median expression level and expression noise for the genotypes surveyed, (ii) Supplementary Tables, including Table S1 (Position and nature of polymorphisms in the *TDH3* promoter) and Table S2 (List of oligonucleotides used to construct strains), Supplementary File 1 (Input tables for analyses of flow cytometry data), Supplementary File 2 (Output tables including all expression and fitness data), and Supplementary File 3 (R scripts used to analyze flow cytometry data).

## Acknowledgements

We thank Chetna Gopinath for technical assistance, Brian Metzger, David Yuan, Bing Yang, and Andrea Hodgins-Davis for helpful discussions, and Brian Metzger, Jennifer Lachowiec, and Andrea Hodgins-Davis for comments on the manuscript. This work was supported by a European Molecular Biology Organization postdoctoral fellowship (EMBO ALTF 1114-2012) to F.D., a Ph.D. fellowship from Ecole Doctorale BMIC de Lyon to W.T., and grants from the National Science Foundation (MCB-1021398) and National Institutes of Health (R01GM108826 and 1R35GM118073) to P.J.W.

## Supplementary Materials for Duveau et al.: “Fitness effects of *cis*-regulatory variants in the Saccharomyces cerevisiae TDH3 promoter”

### Materials and Methods

*Strains used to quantify TDH3 promoter activity*

The distributions of effects of mutations and polymorphisms on *P_TDH3_* activity were described in (Metzger et al. 2015). In this previous study, 236 strains of *S. cerevisiae* were made in which each contained one of the 241 G:C to A:T base pair transitions in the 678bp *TDH3* promoter (*P_TDH3_*) induced by site directed mutagenesis. These *P_TDH3_* alleles were used to drive expression of a yellow fluorescent protein (YFP), with the entire *P_TDH3_-YFP* reporter gene inserted near the *SWH1* pseudogene on chromosome 1 at position 199,270 in the genetic background of the haploid *MAT**a*** strain BY4724 (Brachmann et al. 1998). One of these strains, m131, was excluded from this study because it appeared to have acquired a secondary mutation that affected fluorescence since data were collected for Metzger et al. (2015). Locations of the remaining 235 mutations are described in the Mutation. Positions worksheet in Supplementary File 1. 27 *P_TDH3_* haplotypes observed among 85 strains of *S. cerevisiae*, containing a total of 45 different polymorphisms, were introduced upstream of the same reporter gene used to assess the effects of the point mutations. The evolutionary history of these *P_TDH3_* haplotypes was inferred by comparing them to each other and to the homologous sequence from 15 distantly related strains of *S. cerevisiae* and from all *Saccharomyces sensu stricto* species. For pairs of haplotypes that differed by a single polymorphism, the effect of that polymorphism was inferred by comparing activity of the two haplotypes. For natural haplotypes that differed by two polymorphisms from their closest neighbor in the haplotype network, the effects of individual polymorphisms were determined by constructing intermediate haplotypes connecting natural haplotypes to their closest neighbor through single mutational steps. Natural haplotypes differing by more than two polymorphisms from all other haplotypes were excluded, reducing the number of unique polymorphisms from 45 to 30. In seven instances, the order of occurrence of two consecutive polymorphisms remained unresolved. In these cases, we calculated the effects of the two polymorphisms according to the two possible mutational paths (ab -> Ab -> AB and ab -> aB -> AB). A linear model was then applied to test for the significance of the interaction between the genotypes at the two polymorphic sites on fluorescence levels. Among the seven pairs of polymorphisms tested, none of the interaction terms was significant (*P* > 0.05) after Bonferroni correction for multiple testing, although significant epistasis was detected for two pairs of polymorphisms without applying the multiple test correction (*P* = 0.02 and *P* = 0.008). We therefore averaged the effects of the polymorphisms quantified on the two possible mutational paths. Overall, this approach allowed us to estimate the effect of 30 individual polymorphisms in the promoter context where they most likely arose. A summary of variants present in each of the natural and inferred haplotypes is shown in Table S1, and the 45 pairs of haplotypes compared to infer the effects of 30 polymorphisms are shown in the Haplotype. Network worksheet of Supplementary File 1.

The 14 *P_TDH3_* genotypes used to establish the relationship between TDH3 expression level and fitness are described in the Expression.vs.Fitness worksheet in Supplementary File 2. These genotypes were selected to cover a broad and homogenous range of promoter activity ranging from 0% to over 100%. To decrease the activity of the promoter, eight genotypes were chosen from the 235 strains constructed in Metzger et al. (2015) with single mutations in *P_TDH3_* (Figure 1A). The un-mutated progenitor strain of these lines, defined as the wild type allele of *P_TDH3_*, was also included as the reference allele. As described in Metzger et al. (2015), our reference allele differs from the S288c *P_TDH3_* sequence by a single base (G293a). The derived nucleotide is present in the 235 mutant alleles of *P_TDH3_*, while the ancestral nucleotide is present in all natural and intermediate haplotypes. Because this single nucleotide difference was not found to interact epistatically with other mutations (Metzger et al. 2015), our relative measures of fluorescence (see below) correct for its effect on expression level. The original strain BY4724, which did not contain the *P_TDH3_-YFP* reporter gene, was used to determine the baseline level of auto-fluorescence in cells when the promoter activity was effectively 0%. To increase expression of the YFP reporter gene, we created three strains with a duplication of the entire *P_TDH3_-YFP* construct (Figure 1A). One of these strains contained two copies of the wild type (S288c) *TDH3* promoter, while the other two strains carried a single mutation in each copy of the promoter (see Expression.vs.Fitness worksheet in Supplementary File 2). These duplications were created in a strain described in Metzger et al. (2016) that contains a few genetic differences from the strains characterized in Metzger et al. (2015). This haploid *MATα* strain contains alleles of *SAL1*, *CAT5* and *MIP1* that decrease the penetrance of the petite phenotype (Dimitrov et al. 2009), alleles of *RME1* and *TAO3* that increase sporulation efficiency in diploids (Deutschbauer and Davis 2005), and has the *P_TDH3_-YFP* construct inserted at the *HO* locus. We previously showed that mutations in *P_TDH3_* had similar effects on the expression of the reporter gene in the two genetic backgrounds (Metzger et al. 2016). To create the different duplications, we first amplified by PCR each allele of the *P_TDH3_-YFP* reporter gene with oligonucleotides 2687 and 1893 (Table S2) as well as the *URA3* cassette from the *pCORE-UH* plasmid (Stuckey et al. 2011) with oligonucleotides 2688 and 2686 (Table S2). Next, *URA3-P_TDH3_-YFP* fusions were obtained using overlap extension PCR. These fusions were amplified by PCR with oligonucleotides 2684 and 2683 (Table S2) containing homology arms for insertion downstream of *P_TDH3_-YFP* in strains carrying the reporter gene inserted at *HO* and with desired mutations in *P_TDH3_* (Metzger et al. 2016). Therefore, the resulting strains carried a *P_TDH3_-YFP-URA3-P_TDH3_-YFP* duplicate reporter gene with the same allele of *P_TDH3_* twice. For each strain, Sanger sequencing confirmed the correct sequence of the entire construct. Finally, to control for the effect of *URA3* on fluorescence level, we inserted the *URA3* cassette amplified with primers 2684 and 2685 (Table S2) downstream of *P_TDH3_-YFP* in a strain carrying the wild type allele of *P_TDH3_* at *HO*.

#### Quantifying P_TDH3_ activity by flow cytometry

For each strain, the median fluorescence level (and the standard deviation of fluorescence level) of at least 1500 cells was quantified by flow cytometry and used as a measure of the activity of the *P_TDH3_* allele driving expression of *YFP* in that strain. The activity of the 235 *P_TDH3_* alleles with single G:C to A:T transitions and the natural and intermediate haplotypes was previously assessed in nine replicate populations upon growth to saturation in rich medium (YPD) containing 10 g/L of yeast extract, 20 g/L of peptone and 20 g/L of dextrose (Metzger et al. 2015). We used here a similar approach to quantify the fluorescence levels of the three strains carrying duplication of the reporter gene and of the reference strain containing a single copy of the wild type reporter gene and the *URA3* marker. Each strain was revived from a frozen glycerol stock for 48 hours on medium lacking uracil (5 g/L ammonium sulfate, 1.7 g/L yeast nitrogen base, 1.44 g/L CSM-Ura, 3% v/v glycerol, 20 g/L agar) and then arrayed in eight replicate populations in 96-well plates containing 0.5 mL of YPD per well. After 22 hours of growth at 30**°**C, samples underwent three rounds of dilution to 2.5 x 10^4^ cells/mL in fresh YPD followed by 12 hours of growth at 30**°**C. Then, cultures were diluted to 2.5 x 10^6^ cells/mL in 96-well plates filled with 0.5 mL PBS per well. For each sample, the size and fluorescence level of more than 1000 individual cells were quantified using a BD Accuri C6 flow cytometer coupled with a HyperCyt autosampler. A 530/30 nm optical filter was used to acquire the fluorescence emitted by YFP. This new dataset and the data collected in (Metzger et al. 2015) were all analyzed using custom R scripts (Supplementary File 3) similar to Metzger et al (2016). Briefly, events that did not correspond to single cells were filtered out using the clustering functions of the Bioconductor package *flowClust*. The relationship between forward scatter area (FSC.A) and forward scatter height (FSC.H) was then used to distinguish single cells, budding cells and pairs of cells. Non-fluorescent debris and dead cells were then removed, retaining only events corresponding to healthy fluorescent cells. The next step was to calculate a fluorescence level that did not depend on cell size. Indeed, the fluorescence intensity FL1.A is strongly correlated with FSC.A. To remove this correlation, we applied a principal component analysis on the logarithm of FSC.A and the logarithm of FL1.A for all filtered events. We then transformed the data following a rotation centered on the intersection between the two eigenvectors and of an angle defined by the first eigenvector and the vector between the origin and the intersection between the two eigenvectors. Transformed values of FL1.A were then divided by transformed values of FSC.A, yielding single cell fluorescence levels uncorrelated with FSC.A. For each sample, we then calculated the median fluorescence level of all cells and subtracted the auto-fluorescence measured from the strain without the reporter gene. The fluorescence signal of each sample was divided by the average fluorescence signal of the reference strain carrying the wild type allele of *P_TDH3_* and averaged across replicates to obtain the relative fluorescence levels shown in Figure 1A and Figure 2C. We also calculated the noise of fluorescence levels (*i.e*. the variability of expression level among genetically identical cells) for each sample by dividing the standard deviation of fluorescence levels across all cells in the sample by the median fluorescence level after correction for auto-fluorescence. Relative noise shown in Figure S1 of the Supplementary Text was calculated by dividing the noise of each sample by the average noise measured for the strain carrying the wild type allele of *P_TDH3_* and by averaging the result across replicate populations.

#### Strains included in the fitness assays

To assess how changes in TDH3 expression level affected fitness, we introduced each of the 14 alleles of the *TDH3* promoter described in the Expression.vs.Fitness worksheet in Supplementary File 2 at the native *TDH3* locus (Figure 1B). First, we created strain YPW1121 by replacing the native *TDH3* promoter with a *CORE-UK* cassette (Stuckey et al. 2011) amplified using oligonucleotides 1909 and 1910 (Table S2) in the *MAT***a** strain YPW1001 carrying the wild type allele of the *P_TDH3_-YFP* reporter gene at *HO* (as well as a *NatMX4* drug resistance marker). YPW1001 genome also contained the alleles of *SAL1*, *CAT5* and *MIP1* decreasing petite frequency (Dimitrov et al. 2009) and the alleles of *RME1* and *TAO3* increasing sporulation efficiency (Deutschbauer and Davis 2005). For the non-duplicated alleles, we used genomic DNA from strains used in the fluorescence assays to amplify *P_TDH3_* alleles located upstream of *YFP* with oligonucleotides 2425 and 1305 (Table S2) specific to the insertion site on chromosome I and to the *YFP* coding sequence (to avoid amplification of the native *TDH3* promoter). The resulting PCR products were then amplified with 90 bp oligonucleotides 1914 and 1900 (Table S2) including 70 nucleotides of homology to the native *TDH3* locus at their 5’ end. These products were then transformed in strain YPW1121 to replace the *CORE-UK* cassette with the mutated alleles of *P_TDH3_*. Transformants were selected on 5-FOA plates and successful insertions of *P_TDH3_* alleles were confirmed by Sanger sequencing. To determine fitness when *TDH3* expression level was 0%, we generated strain YPW1177 where the entire *TDH3* promoter and coding sequence were deleted. This was done by transforming strain YPW1121 with a PCR product obtained with primers 1345 and 1962 (Table S2) including homology to the sequences immediately upstream of the *TDH3* promoter and downstream of the *TDH3* stop codon. To quantify the fitness effects of increased *TDH3* expression level, we created three strains with duplication of the entire *TDH3* gene. In these strains, the alleles of the two copies of *P_TDH3_* at the native locus correspond to the *P_TDH3_* alleles present in the three strains with duplicated reporter genes (described in Expression.vs.Fitness worksheet in Supplementary File 2). First, we amplified with oligonucleotides 2687 and 1893 (Table S2) the *TDH3* promoter and coding sequence from the strains carrying at the native *TDH3* locus the three alleles to be duplicated. Then, the *URA3* cassette amplified from the *pCORE-UK* plasmid (Stuckey et al. 2011) with oligonucleotides 2688 and 2686 (Table S2) was fused to each *TDH3* amplicon by overlap extension PCR. The fusion products were amplified using oligonucleotides 2696 and 2693 (Table S2) with homology to *TDH3* 5’UTR and transformed in the genome of the three strains carrying the corresponding alleles of *P_TDH3_* at the native locus. As a result, the structure of the *TDH3* locus after duplication was *P_TDH3_-TDH3-URA3-P_TDH3_-TDH3* with both copies of *P_TDH3_* presenting the same allele. Correct sequences of the duplicated genes were confirmed by Sanger sequencing. Lastly, a strain was constructed by inserting the *URA3* cassette amplified with primers 2696 and 2697 (Table S2) downstream of *TDH3* in YPW1001 genome. This strain served as the wild type reference for measuring the fitness consequence of *TDH3* duplications by controlling for the impact of *URA3* expression on fitness. To measure the fitness of the 14 strains described above, each of them was grown for several generations in competition with the same competitor strain YPW1160. This strain expressed a *GFP* reporter gene so that it could be easily distinguished from the 14 strains expressing *YFP* using flow cytometry, as described below. To generate strain YPW1160, the *GFP* coding sequence amplified using oligonucleotides 601 and 2049 (Table S2) was fused to the *KanMX4* drug resistance cassette amplified using oligonucleotides 2050 and 1890 (Table S2) using overlap extension PCR. The PCR product was then amplified with primers 601 and 1890 (Table S2) and transformed into strain YPW1001 to swap the *YFP-NatMX4* cassette at *HO* with *GFP-KanMX4*.

#### Quantifying fitness through competition assays

To precisely quantify the fitness of the 14 strains expressing *TDH3* under different alleles of the promoter, we used highly sensitive competition assays (Figure 2A). First, starting from - 80**°**C glycerol stocks, uracil auxotroph strains were revived on YPG plates (10 g/L yeast extract, 20 g/L peptone, 3% v/v glycerol, 20 g/L agar) and prototroph strains expressing URA3 were revived on plates lacking uracil (5 g/L ammonium sulfate, 1.7 g/L yeast nitrogen base, 1.44 g/L CSM-Ura, 3% v/v glycerol, 20 g/L agar). Then, the 10 [ura-] strains were arrayed in six 96-well plates containing 0.5 mL of YPD per well. Each plate contained 24 replicates of the reference strain, 24 replicates of a strain carrying a strong effect mutation in *P_TDH3_* and 24 replicates of two other mutant strains (Layout.Fitness worksheet in Supplementary File 1). The four strains expressing URA3 (three strains with *TDH3* duplication and the corresponding reference strain) were arrayed in eight replicates on a separate plate that also contained other strains not included in this study (Layout.Fitness worksheet in Supplementary File 1). The fitness of these strains was measured in three separate experiments, with each experiment including a subset of the seven plates described above (Layout.Expression worksheet in Supplementary File 1). We also inoculated a matching set of 96-well plates for each experiment with only the GFP-expressing strain YPW1160 in every well. Plates were grown for 22 hours at 30**°**C on a roller drum to maintain cells in suspension. After growth, the optical density at 620 nm of all samples was recorded using a Sunrise plate reader, allowing us to calculate the average cell density for each plate. Using this information, we next mixed for each sample an approximately equal number of cells expressing YFP ([YFP+]) and of cells expressing GFP ([GFP+]) in clean plates. Then, an average of 4 x 10^4^ mixed cells were diluted in 0.45 mL of fresh YPD medium so that the average cell density on each plate would reach approximately 5 x 10^6^ cells/mL after 12 hours of growth at 30**°**C. After this initial phase of competitive growth, we performed two more rounds of cell density measurement, dilution to fresh YPD and growth for 12 hours so that all samples acclimated to constant growth in glucose. After this acclimation phase, the procedure of cell density measurement, dilution and growth was repeated three more times, except that samples were grown for 10 hours between each dilution. After each dilution and also after the last round of growth, culture leftovers were diluted to 1.5 x 10^6^ cells/mL in 0.3 mL of PBS in 96-well plates, placed on ice and run on the BD Accuri C6 flow cytometer. A 488 nm laser was used for excitation and the fluorescence emitted was acquired through two different optical filters at 510/15 nm and 585/40 nm, allowing separation of the GFP and YFP signals. At least 5,000 events were recorded for each sample at each of the four time points separated by 10 hours or ~7 generations of competitive growth (for a total of ~21 generations). The number of generations was determined for each plate based on the changes in cell density observed before and after the 10 hours of growth. Cell densities were estimated from the optical densities of the wild type strains present on each plate.

We then calculated the fitness of each population of [YFP+] cells based on the evolution of the ratio of [YFP+] by [GFP+] cells over time. To determine the frequency of [YFP+] and [GFP+] cells in each sample, we analyzed the flow cytometry data using custom R scripts (Supplementary File 3). After logarithmic transformation of raw data, we filtered out artifacts with extreme values of forward scatter (FSC.A and FSC.H) or fluorescence intensity (FL1.H acquired through the 510/15 filter and FL2.H acquired through the 585/40 filter). Three populations could be distinguished based on FL2.H/FL1.H ratio: events with high FL2.H/FL1.H values corresponded to [YFP+] cells, events with low FL2.H/FL1.H values corresponded to [GFP+] cells and the smaller population of events with intermediate values of FL2.H/FL1.H corresponded to doublets of one [YFP+] cell and one [GFP+] cell scored simultaneously (these events were discarded). To separate [YFP+] and [GFP+] populations, we used dynamic gates that could accommodate for shifts in fluorescence levels instead of using arbitrary gates. We first performed a principal component analyses on FL1.H and FL2.H data. Then, we fitted a density function using Kernell estimates to the second principal component scores and we separated [YFP+] and [GFP+] populations using two thresholds corresponding to the local minima of the density function. Using these thresholds, we then counted the number of [YFP+] and [GFP+] cells analyzed by flow cytometry for each sample. Next, we calculated for each plate the number of generations of the reference strain between the four time points based on cell densities measured for the 24 wild type replicates at each time point and based on dilution factors. Finally, for each sample the number of generations calculated for the plate was regressed on the logarithm of the ratio of the number of [YFP+] cells divided by the number of [GFP+] cells using the R function *lm*. The exponential of the regression coefficient corresponded to the fitness of the [YFP+] population relative to the [GFP+] population (Hartl & Clark, 2007). We then divided the fitness of the 96 samples on each plate by the mean fitness of the 24 replicates of the reference strain on that plate, resulting in a measure of fitness expressed relative to the fitness of the reference strain (without mutation in the *TDH3* promoter). We used the *t.test* function in R for statistical comparison of the mean fitness of each strain to the mean fitness of the reference strain replicates present on the same plate.

#### Relationship between P_TDH3_ activity and fitness

To characterize the relationship between *TDH3* expression level and fitness, we performed a local regression (LOESS regression) of the mean fluorescence levels measured for the 14 strains expressing *YFP* under different alleles of *P_TDH3_* on the mean fitness measured for the 14 strains expressing *TDH3* under the same *P_TDH3_* alleles (Figure 2C). We used the R function *loess* with a span of 0.8 to control the degree of smoothing.

#### Access to data and analysis scripts

Flow cytometry data used in this work is available through the Flow Repository (https://flowrepository.org): Repository ID = FR-FCM-ZZBN for data from Metzger et al. (2015) used to quantify the effects of 235 mutations and 30 polymorphisms in *P_TDH3_* on expression, Repository ID = FR-FCM-ZY8Y for data collected here to quantify the effects of duplications of the reporter gene on expression and Repository ID = FR-FCM-ZY7E for data collected here to quantify the fitness effects of 14 alleles of *P_TDH3_*. The three R scripts used to perform the analyses described above (“Analysis of flow data for quantifying average fluorescence levels”, “Analysis of flow data for quantifying fitness” and “Plot data shown on Figure 1”) have been concatenated and provided as Supplementary File 3. Data tables used by these scripts are provided in Supplementary File 1. Specifically, Supplementary File 1 includes a description of each 96-well plate used to analyze expression (Layout.Expression.txt worksheet), a summary of the type of mutation (Mutation.Types.txt worksheet) and position of mutation (Mutation.Positions.txt worksheet) in each strain, a list of haplotypes compared to infer the effects of polymorphisms (Haplotype.Network.txt worksheet), a description of each 96-well plate used to analyze fitness (Layout.Fitness.txt worksheet), and measures used to convert optical density to cell density during fitness assays (Optical.Density.txt worksheet). The FCS files needed for each analysis must be placed in the same parent directory and must appear in the same order as in Layout.Expression.txt and Layout.Fitness.txt worksheets. Supplementary File 2 includes summary tables of expression and fitness for each genotype: a summary of strains, mutations, expression, and fitness used to plot the fitness function shown in Figure 2C (Expression.vs.Fitness worksheet), a summary of estimated expression level for the 235 mutations (Expression.Mutations worksheet), 45 haplotypes of *P_TDH3_* (Expression.Haplotypes worksheet), 30 individual polymorphisms (Expression.Polymorphisms worksheet), and summaries of expression (Summary.Expression worksheet) and fitness (Summary.Fitness worksheet) in the format used by the script in Supplementary File 3 to produce Figure 1.

---

**Supplementary Figure 1.**
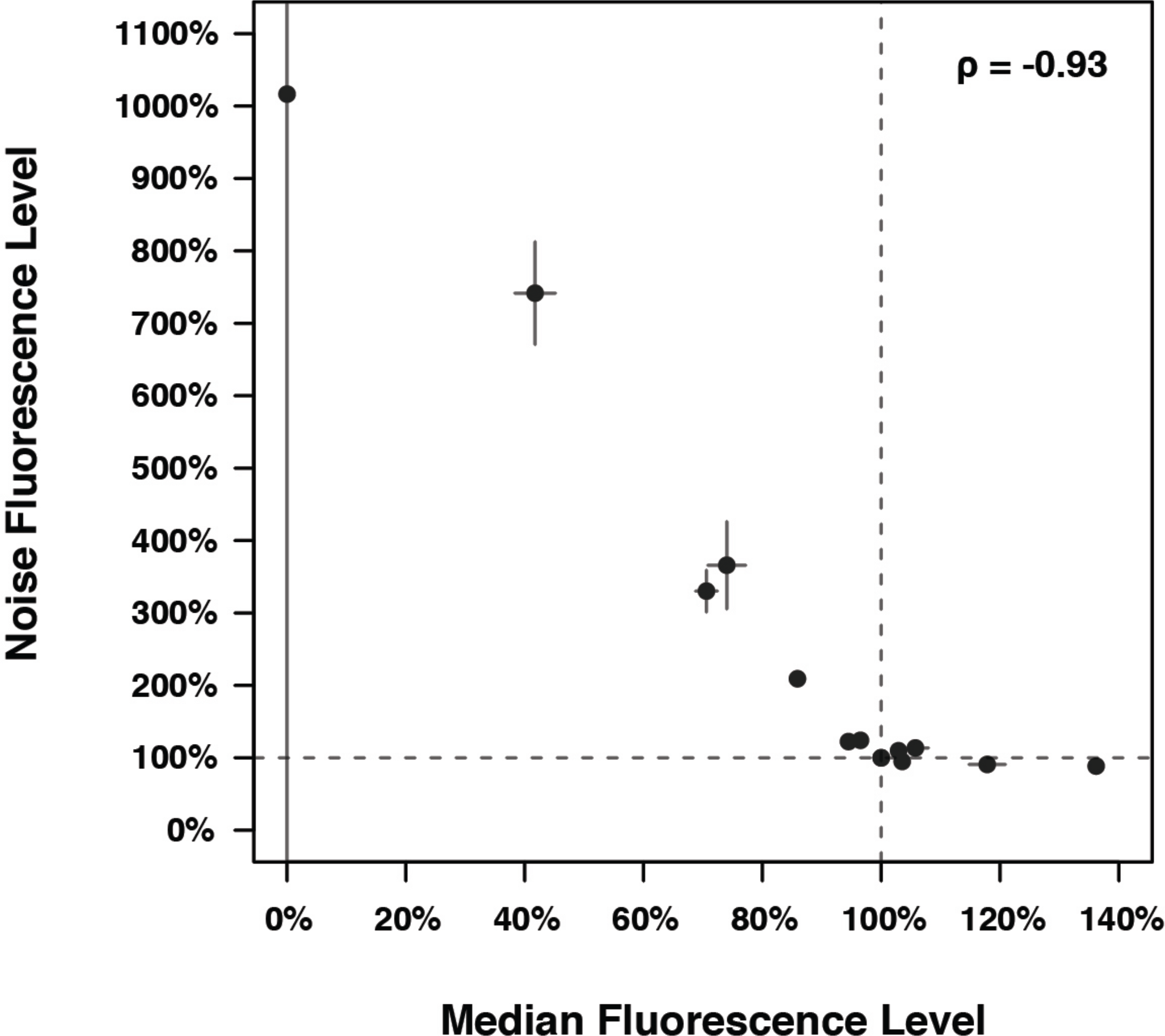
Effects of *P_TDH3_* mutations on expression level and expression noise are strongly correlated. For each of the *P_TDH3_-YFP* reporter genes assayed, the median and noise of fluorescence averaged across replicates is shown. Median fluorescence levels are expressed relative to the fluorescence level of the wild type allele and noise was calculated as the standard deviation of fluorescence among individual cells divided by the median fluorescence level, and it was also expressed relative to the noise of the wild type allele. Error bars represent 95% confidence intervals, and Spearman’s rho was used to assess the sign and strength of the correlation.

## Supplementary Tables and Files

Supplementary Tables, including Table S1 (Position and nature of polymorphisms in the *TDH3* promoter) and Table S2 (List of oligonucleotides used to construct strains).

Supplementary File 1. Input tables for analyses of flow cytometry data.

Supplementary File 2. Output tables including all expression and fitness data.

Supplementary File 3. R scripts used to analyze flow cytometry data.

## References

Alvarez M, Schrey AW, Richards CL. 2015. Ten years of transcriptomics in wild populations: what have we learned about their ecology and evolution? Mol. Ecol. 24:710–725.

Bergen AC, Olsen GM, Fay JC. 2016. Divergent *MLS1* Promoters Lie on a Fitness Plateau for Gene Expression. Mol. Biol. Evol. 33:1270–1279.

Branco P, Francisco D, Chambon C, Hébraud M, Arneborg N, Almeida MG, Caldeira J, Albergaria H. 2014. Identification of novel GAPDH-derived antimicrobial peptides secreted by Saccharomyces cerevisiae and involved in wine microbial interactions. Appl. Microbiol. Biotechnol. 98:843–853.

Carroll SB. 2008. Evo-devo and an expanding evolutionary synthesis: a genetic theory of morphological evolution. Cell 134:25–36.

Deutschbauer AM, Jaramillo DF, Proctor M, Kumm J, Hillenmeyer ME, Davis RW, Nislow C, Giaever G. 2005. Mechanisms of haploinsufficiency revealed by genome-wide profiling in yeast. Genetics 169:1915–1925.

Dykhuizen DE, Dean AM, Hartl DL. 1987. Metabolic flux and fitness. Genetics 115:25–31.

Fay JC, Wittkopp PJ. 2008. Evaluating the role of natural selection in the evolution of gene regulation. Heredity (Edinb) 100:191–199.

Gancedo JM. 1998. Yeast carbon catabolite repression. Microbiol. Mol. Biol. Rev. 62:334–361.

Gilad Y, Oshlack A, Rifkin SA. 2006. Natural selection on gene expression. Trends in Genetics 22:456–461.

Kacser H, Burns JA. 1973. The control of flux. Symp. Soc. Exp. Biol. 27:65–104.

Kafri M, Metzl-Raz E, Jona G, Barkai N. 2016. The Cost of Protein Production. Cell Rep 14:22–31.

Keren L, Hausser J, Lotan-Pompan M, Vainberg Slutskin I, Alisar H, Kaminski S, Weinberger A, Alon U, Milo R, Segal E. 2016. Massively Parallel Interrogation of the Effects of Gene Expression Levels on Fitness. Cell 166:1282–1294.e18.

Makanae K, Kintaka R, Makino T, Kitano H, Moriya H. 2013. Identification of dosage-sensitive genes in *Saccharomyces cerevisiae* using the genetic tug-of-war method. Genome Research 23:300–311.

McAlister L, Holland MJ. 1985. Isolation and characterization of yeast strains carrying mutations in the glyceraldehyde-3-phosphate dehydrogenase genes. J. Biol. Chem. 260:15013–15018.

Metzger BPH, Duveau F, Yuan DC, Tryban S, Yang B, Wittkopp PJ. 2016. Contrasting Frequencies and Effects of *cis-* and *trans-*Regulatory Mutations Affecting Gene Expression. Mol. Biol. Evol. 33:1131–1146.

Metzger BPH, Yuan DC, Gruber JD, Duveau F, Wittkopp PJ. 2015. Selection on noise constrains variation in a eukaryotic promoter. Nature 521:344–347.

Newman JRS, Ghaemmaghami S, Ihmels J, Breslow DK, Noble M, DeRisi JL, Weissman JS. 2006. Single-cell proteomic analysis of *S*. *cerevisiae* reveals the architecture of biological noise. Nature 441:840–846.

Perfeito L, Ghozzi S, Berg J, Schnetz K, Lässig M. 2011. Nonlinear fitness landscape of a molecular pathway. PLoS Genet. 7:e1002160.

Ranz JM, Machado CA. 2006. Uncovering evolutionary patterns of gene expression using microarrays. Trends in Ecology & Evolution 21:29–37.

Rest JS, Morales CM, Waldron JB, Opulente DA, Fisher J, Moon S, Bullaughey K, Carey LB, Dedousis D. 2013. Nonlinear fitness consequences of variation in expression level of a eukaryotic gene. Mol. Biol. Evol. 30:448–456.

Rich MS, Payen C, Rubin AF, Ong GT, Sanchez MR, Yachie N, Dunham MJ, Fields. 2016. Comprehensive Analysis of the *SUL1* Promoter of *Saccharomyces cerevisiae*. Genetics. 203, 191–202.

Ringel AE, Ryznar R, Picariello H, Huang K-L, Lazarus AG, Holmes SG. 2013. Yeast Tdh3 (glyceraldehyde 3-phosphate dehydrogenase) is a Sir2-interacting factor that regulates transcriptional silencing and rDNA recombination. PLoS Genet. 9:e1003871.

Wang Z, Zhang J. 2011. Impact of gene expression noise on organismal fitness and the efficacy of natural selection. Proc. Natl. Acad. Sci. U.S.A. 108:E67–E76.

Wittkopp PJ, Kalay G. 2012. *Cis*-regulatory elements: molecular mechanisms and evolutionary processes underlying divergence. Nat Rev Genet 13:59–69.

Wray GA. 2007. The evolutionary significance of *cis*-regulatory mutations. Nat Rev Genet 8:206–216.

## References

Brachmann CB, Davies A, Cost GJ, Caputo E, Li J, Hieter P, Boeke JD. 1998. Designer deletion strains derived from Saccharomyces cerevisiae S288C: a useful set of strains and plasmids for PCR-mediated gene disruption and other applications. Yeast 14:115–132.

Deutschbauer AM, Davis RW. 2005. Quantitative trait loci mapped to single-nucleotide resolution in yeast. Nat. Genet. 37:1333–1340.

Dimitrov LN, Brem RB, Kruglyak L, Gottschling DE. 2009. Polymorphisms in multiple genes contribute to the spontaneous mitochondrial genome instability of Saccharomyces cerevisiae S288C strains. Genetics 183:365–383.

Hartl DL, Clark AG. 2007. Principles of Population Genetics. 4^th^ Edition. Sinauer Associates.

Metzger BPH, Duveau F, Yuan DC, Tryban S, Yang B, Wittkopp PJ. 2016. Contrasting Frequencies and Effects of cis- and trans-Regulatory Mutations Affecting Gene Expression. Mol. Biol. Evol. 33:1131–1146.

Stuckey S, Mukherjee K, Storici F. 2011. In vivo site-specific mutagenesis and gene collage using the delitto perfetto system in yeast Saccharomyces cerevisiae. Methods Mol. Biol. 745:173–191.

